# Learning probabilistic representations with randomly connected neural circuits

**DOI:** 10.1101/478545

**Authors:** Ori Maoz, Gašper Tkacčik, Mohamad Saleh Esteki, Roozbeh Kiani, Elad Schneidman

## Abstract

The brain represents and reasons probabilistically about complex stimuli and motor actions using a noisy, spike-based neural code. A key building block for such neural computations, as well as the basis for supervised and unsupervised learning, is the ability to estimate the surprise or likelihood of incoming high-dimensional neural activity patterns. Despite progress in statistical modeling of neural responses and deep learning, current approaches either do not scale to large neural populations or cannot be implemented using biologically realistic mechanisms. Inspired by the sparse and random connectivity of real neuronal circuits, we present a new model for neural codes that accurately estimates the likelihood of individual spiking patterns and has a straightforward, scalable, efficiently learnable, and realistic neural implementation. This model’s performance on simultaneously recorded spiking activity of >100 neurons in the monkey visual and prefrontal cortices is comparable or better than that of current models. Importantly, the model can be learned using a small number of samples, and using a local learning rule that utilizes noise intrinsic to neural circuits. Slower, structural changes in random connectivity, consistent with rewiring and pruning processes, further improve the efficiency and sparseness of the resulting neural representations. Our results merge insights from neuroanatomy, machine learning, and theoretical neuroscience to suggest random sparse connectivity as a key design principle for neuronal computation.

---

The majority of neurons in the central nervous system know about the external world only by observing the activity of other neurons. Neural circuits must therefore learn to represent information and reason based on the regularities and structure in spiking patterns coming from upstream neurons, in a largely unsupervised manner. Since the mapping from stimuli to neural responses (and back) is probabilistic [1, 2, 3], and the spaces of stimuli and responses are exponentially large, neural circuits must be performing a form of statistical inference by generalizing from the previously observed spiking patterns [4, 5, 6, 7]. Nevertheless, circuit mechanisms that implement such probabilistic computations remain largely unknown.

A biologically plausible neural architecture that would allow for such probabilistic computations would ideally be scalable and could be trained by a local learning rule in an unsupervised fashion. Current approaches satisfy some, but not all, of the above properties. Top-down approaches suggest biologically plausible circuits that solve particular computational tasks, but often rely on explicit “teaching signals” or do not even specify how learning could take place [8, 9, 10, 11, 12, 13, 14]. Notably, an architecture designed for a particular task will typically not support other computations, as done in the brain. Lastly, top-down models relate to neural data on a qualitative level, falling short of reproducing the detailed statistical structure of neural activity across large neural populations. In contrast, bottom-up approaches grounded in probabilistic modeling, statistical physics, or deep neural networks, can yield concise and accurate models of the joint activity of the neural population in an unsupervised fashion [15, 16, 17, 18, 19, 20, 21, 22, 23, 24, 25, 26, 27]. Unfortunately, these models are difficult to relate to the mechanistic aspects of neural circuit operation or computation, because they use architectures and learning rules that are non-biological or non-scalable.

A neural circuit that would learn to estimate the probability of its inputs would merge these two approaches: rather than implementing particular tasks or extracting specific stimulus features, computing the likelihood of the input gives a universal ‘currency’ for the neural computation of different circuits. Such circuit could be used and reused by the brain as a recurring motif, in a modular and hierarchical manner for a variety of sensory, motor, and cognitive contexts, including for feature learning. This would remove the need for many specialized circuits for different computations. Consequently, It would facilitate the adoption of new functions by existing brain circuitry and may serve as an evolutionary principle for creating new modules that communicate and interact with the old ones.

Here we present a simple and highly flexible neural architecture based on spiking neurons, that can efficiently estimate the surprise of its own inputs, thus generalizing from input history in an assumption-free and parsimonious way. This feed-forward circuit can be viewed as implementing a probabilistic model over its inputs, where the surprise of its current input is explicitly represented as the membrane potential of an output (readout) neuron. The circuit is trained by adjusting the connections leading into the output neuron from a set of intermediate neurons, which serve as detectors of random features of the circuit’s input. Unlike many models of neuronal networks, this model relies on local learning in a shallow network, and yet it provides superior performance to state-of-the-art algorithms in estimating the probability of individual activity patterns for large real neural populations. Furthermore, the synaptic connections in the model are learnable with a rule that is biologically plausible and resolves the credit assignment problem [28], suggesting a possible general principle of probabilistic learning in the nervous system.

We consider the joint activity of large groups of neurons recorded from the visual and prefrontal cortices of macaques. Fig. 1a shows examples of activity patterns of 169 neurons, discretized into 20 ms time windows, from the prefrontal cortex of an awake behaving monkey at different times during a classification task. Since for such large populations, particular activity patterns would typically not repeat in the course of the experiment or even in the lifetime of an organism, a neural circuit receiving these patterns as inputs must learn the statistical structure in order to generalize to new, previously unseen, patterns. A neural circuit that estimates the surprise associated with observing a pattern would assess how the new pattern conforms with previously observed patterns, thus generalizing from past inputs without making additional assumptions. In mathematical terms, structure in the input patterns implies that some patterns are more likely to appear than others. This can be described in terms of a probability distribution over input patterns 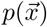, where 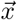is a binary pattern representing the firing (1) or silence (1) of each neuron in the population in a given time bin. The generic notion of surprise of observing an input pattern 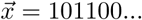 appearing with probability 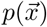 is then given by − 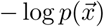 [29].

**Figure 1:**
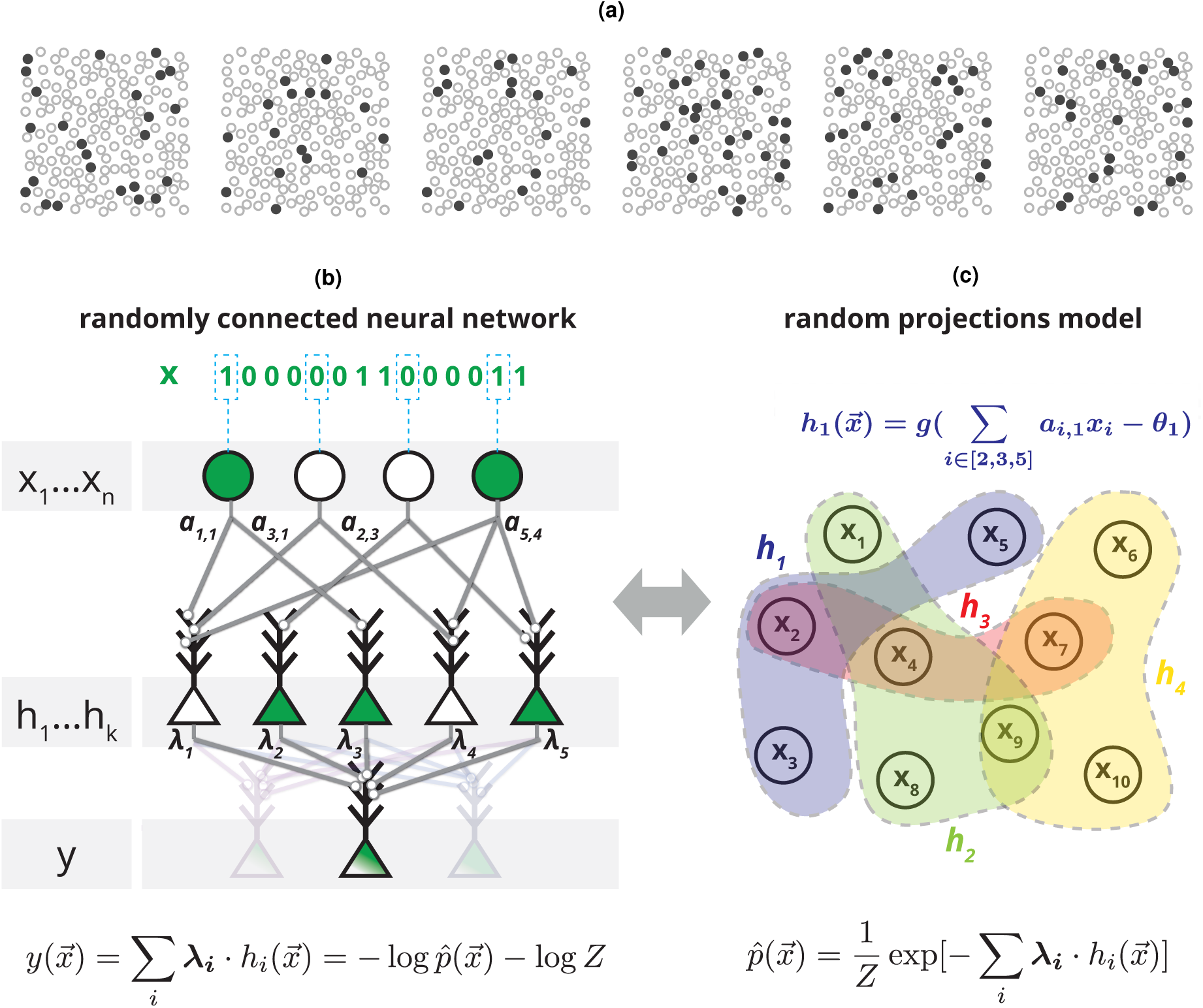
A randomly connected neural network, equivalent to a Random Projections model, that learns to generalize from observed inputs to compute the surprise of novel inputs. **(a)** Examples of six neural population activity patterns at different time points, recorded from 169 neurons in the monkey prefrontal cortex while performing a visual classification task (plotted locations were chosen at random and do not correspond to actual spatial locations). **(b)** Architecture of a random feed-forward neural circuit based on spiking neurons that can learn to respond with the surprise of its input patterns, *x*1*…n*. The input neurons are connected to an intermediate layer of neurons, *h_i_*, with randomly selected synaptic weights *a_ij_*, which then project to an output neuron with synaptic weights *λ_i_*. After learning *λ_i_* the membrane potential of the output neuron 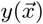 will compute 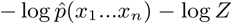, an unnormalized estimate of the surprise, − log *p*(*x*1*…xn*), of the joint input. Note that the same layer of randomly projecting hidden neurons can be reused to simultaneously compute multiple probabilistic models for different output neurons (light color). **(c)** The circuit in (b) is equivalent to a probabilistic model over randomly weighted cliques of neurons, learned by reweighing their contributions, or the maximum entropy model based on random nonlinear statistics of the input.

Fig. 1b illustrates the architecture of a simple and shallow circuit, based on binary neurons, which can learn to respond to input patterns by giving their surprise: The input neurons {*x_j_*} are randomly connected to the neurons in an intermediate layer {*h_i_*}, with randomly selected weights {*a_i,j_*} and so each of the *h_i_*’s computes a non-linear random projection of the input given by *h_i_* = *g*(∑_j_ *a_i,j_x_j_* – *θ_i_*) where *g*() is a threshold function and *θ_i_* is the neuron’s threshold, which we set to a fixed value for all neurons (see SI). These intermediate layer neurons, each serving the role of a feature detector in the input layer, are then connected to a readout neuron, *y*, with weights *λ_i_*. The specific values of *λ_i_*’s thus determine the function that the readout neuron computes based on the projections. The sum of inputs to the readout neuron, or its ‘membrane potential’, is then given by

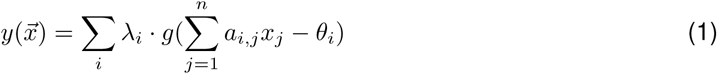

This membrane potential can also be interpreted as 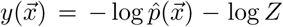, where 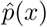 corresponds to an internal model of the inputs:

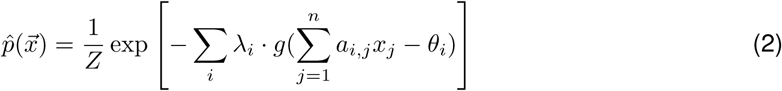

and *Z* is a normalization factor (or partition function). The membrane potential 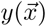 thus reflects an unnormalized internal model of the input distribution or the surprise of the joint inputs, – log *p*(*x_1_*; *x_2_*; …; *x_n_*), up to an additive factor. This factor can be compensated for by learning a bias to the readout neuron’s voltage or its spiking threshold, that would give a normalized value of the surprise (see SI for discussion of possible normalization mechanisms and implementation). We are thus seeking the *λ_i_*’s for which the distribution of inputs 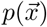 and the internal model 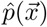 are as similar as possible. Since these are probability distributions, the distance between them is naturally captured by their relative entropy or Kullback-Leibler divergence, and can be minimized by finding the *λ_i_*’s that would maximize the likelihood assigned to inputs by the readout neuron based on its history of input statistics. We recall that Eq. 2 is the well known Boltzmann distribution, offering an alternative interpretation of the function that this circuit computes: given a set of K random functions of the input, *h_i_*’s, find the minimal model that is consistent with the expected values of these functions. This is then the most unstructured description of the data, or the maximum entropy distribution based on the chosen random projections. Yet another interpretation is that this is the reweighting of activity of random cliques or assemblies of neurons [30]. Whichever interpretation one may like, the result is a circuit whose synaptic weights *λ_i_* correspond to the model parameters, and such models can be trained from a set of examples using standard numerical gradient-descent based approaches [31].

The randomly connected neural circuit we described for estimating the surprise is therefore a mechanistic implementation of the probabilistic model based on random projections (RP) described by Fig. 1c (and Eq. 2). Critically, training this RP model requires only changing the synaptic weights *λ_i_* to the output neuron, using a process that requires no extra information about the projections other than that they are sufficiently informative about the input patterns. Thus, the connectivity *a_i,j_* could also be predetermined (evolved) or learned by a separate process (feature selection). This simple design, where the process of selecting the features is distinct from the process of learning how to combine them, sidesteps the well known credit assignment problem where it is unclear which weights are to be updated in each step [28]. Importantly, although the connectivity *a_i,j_* could be optimized or learned by a separate process (more below), purely random connectivity already results in a powerful and flexible probabilistic representation.

The RP model gives an excellent description of the joint activity patterns of large groups of cortical neurons and generalizes from training samples to estimate the likelihood of test data: Figure 2a shows a short segment of spiking patterns of the jointly recorded population activity of 178 neurons from the macaque monkey visual cortex (V1/V2) under anesthesia while moving gratings were presented in the neurons’ receptive fields, and a segment of 169 neurons from the prefrontal cortex while the monkey performed a visual discrimination task. We first evaluated the models on smaller groups of neurons (70 cells from the visual cortex and 50 cells from the prefrontal cortex), where we can directly test the validity of the model because individual activity patterns still repeat. We found that models using 2000 random projections (fit on training data) were highly accurate in predicting the frequency of individual population activity patterns in test data. These populations were strongly correlated as a group, which is reflected by the orders of magnitude prediction errors of the independent model that does not take such correlations into account (left panel). In contrast, maximum entropy models which use pairwise constraints [17, 18, 32] were considerably better (center panel), and RP models were superior with a smaller number of parameters (compared to the pairwise models). For the entire populations of 178 and 169 neurons, where individual activity patterns were so rare that they did not repeat during the experiment, RP models were highly accurate in predicting high-order correlations (SI Fig. S1) and population synchrony in the experimental data (Fig. 2b).

**Figure 2:**
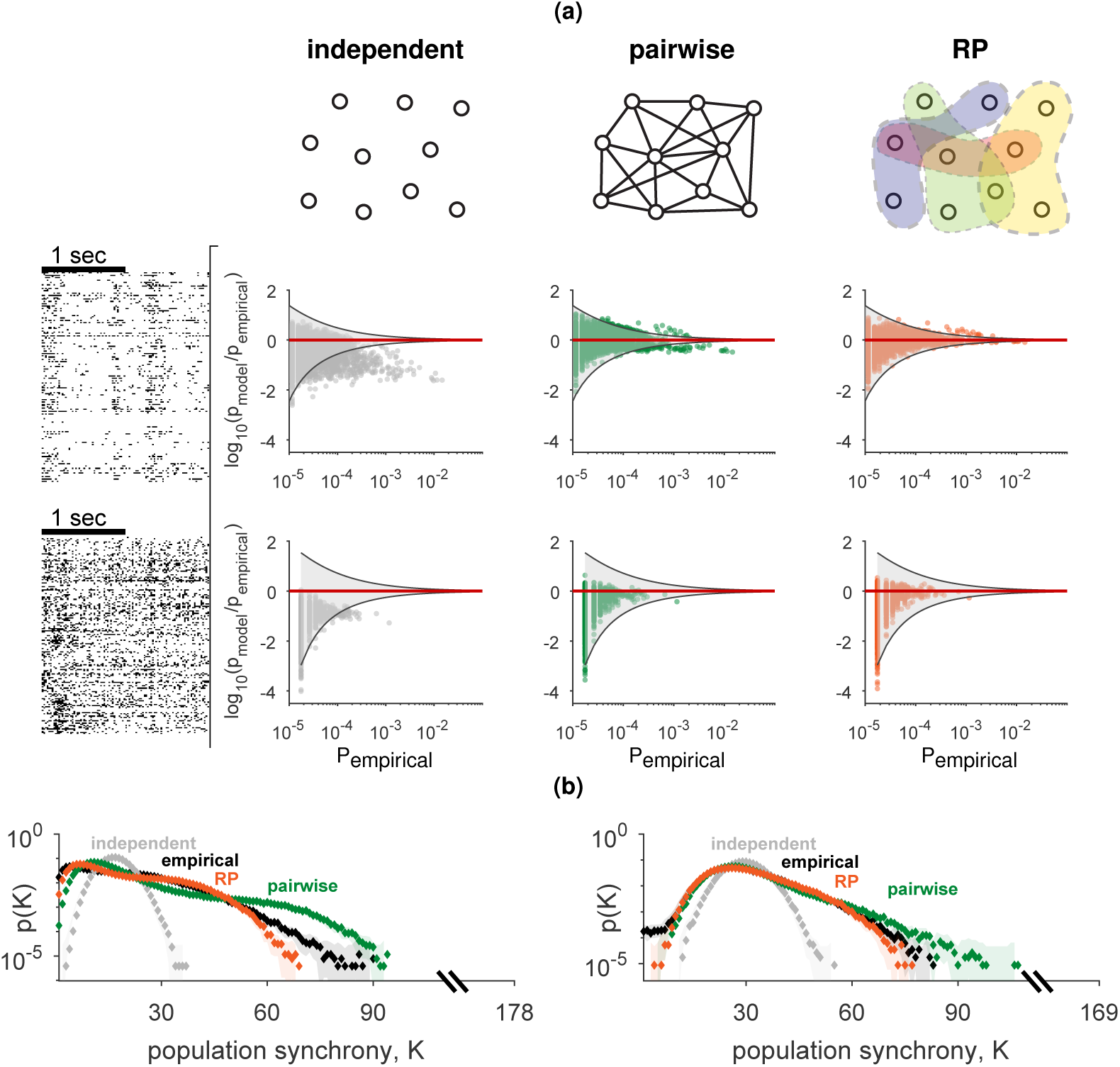
Random Projections models accurately predict the probability of population activity patterns. **(a)** Accuracy of different population models in capturing the frequencies of individual population activity patterns in test data for 70 neurons in the monkey visual cortex (top) or 50 neurons from the monkey prefrontal cortex (bottom): we compare likelihood ratio of models and test data for an independent model (left), pairwise maximum entropy model (middle), and random projection model (right). Grey funnel denotes 99% confidence interval of the likelihood ratio resulting from sampling fluctuations. **(b)** Probability of observing the simultaneous activation of K neurons (population synchrony) in a population of 178 neurons from the primate visual cortex (left) and 169 neurons from the primate prefrontal cortex (right) in test data and model predictions.

Randomly connected circuits have been successfully used in other statistical contexts before: as a design feature in machine learning, specifically for classification [11, 33], or in signal processing for signal reconstruction [34, 35, 36]. Here, in addition to superior performance, random connectivity also allows for greater flexibility of the probabilistic model: since the projections in the model are independent samples of the same class of functions, we can simply add projections (corresponding to adding intermediate neurons in a randomly connected circuit) to improve the accuracy of the model. This allows using as many or as few projections as required, in contrast to pairwise and higher-order correlation based models that are difficult to scale to very large populations [22, 37]. Indeed, the RP models improve monotonically with the number of projections, and become on par with or better than state of the art models, but with less parameters [23], as reflected by both the likelihood of test data of large populations (Fig. 3a, see also SI. Fig. S2c) and direct comparisons in small networks (SI Fig. S2a).

**Figure 3:**
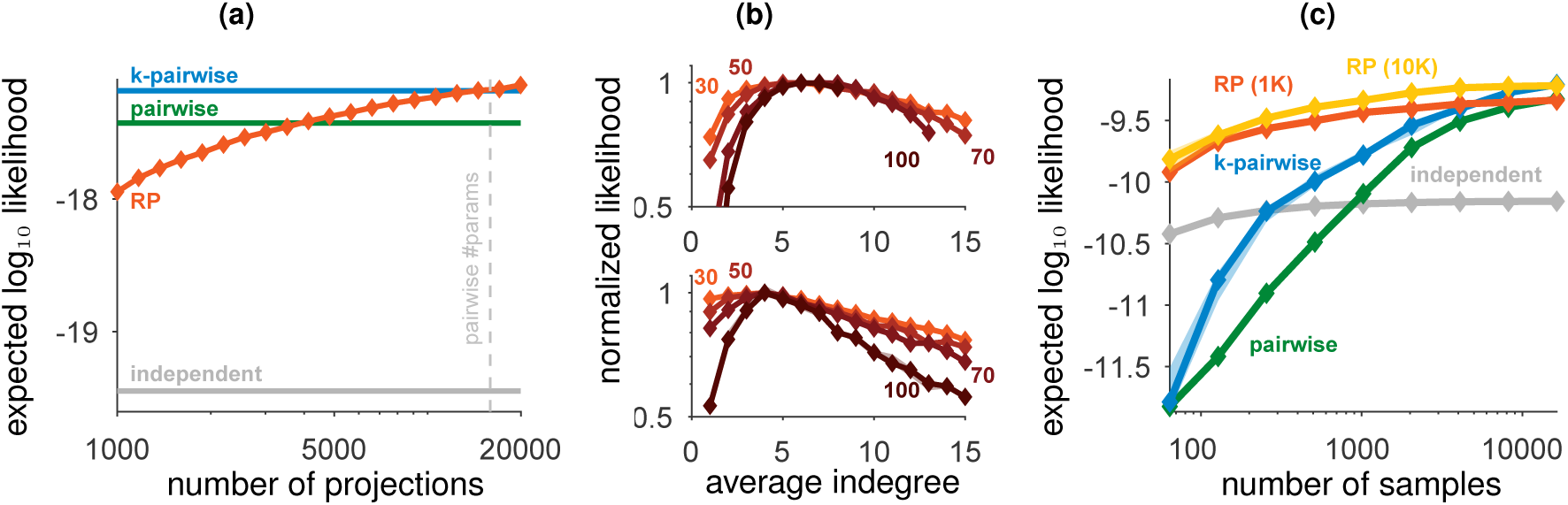
Scalability, optimal sparseness, and efficiency of the Random Projections models. **(a)** Expected likelihood of RP models for held-out data of individual population activity patterns of 178 neurons in the visual cortex as a function of the number of projections used in the model (trained using 100,000 samples); Plotted values are median model performance over random choices of projections and divisions of train/test data (error bars are smaller than the marker size, see SI. Fig. S2c for zoomed-in version). **(b)** Performance of RP models (expected likelihood normalized to a maximum value of 1) with different average indegrees (number of incoming connections) of the intermediate neurons, for the visual cortex (top) and for prefrontal cortex (bottom), each using 131072 input activity patterns. Different curves denote different sizes of input populations, as denoted on each curve. **(c)** Expected likelihood of RP trained on population activity patterns of 100 neurons from the monkey visual cortex as a function of the number of samples in the training data.

The performance of the RP models has very little variance, for different randomly chosen sets of projections (SI Fig. S2b), reflecting that the exact sets of random projections used in each model are unimportant, and can be replaced. Different choices of generating the random projections *a_i,j_* had little effect on the model performance (SI Fig. S3a), and RP models using other classes of random functions we tested were inferior to those using threshold-linear neurons (SI Fig. S3b). We find that for populations of different sizes, RP models were most accurate when the projections were sparse in terms of the number of *a_i,j_* weights that were not zero, corresponding to neural circuits with a low average indegree of their intermediate layer. Thus sparseness, which has been suggested as a design principle for neural computation [38], emerges in the RP models as their optimal operational regime. The optimal average indegree value ranged between ~4 for the prefrontal cortex to ~7 for the visual cortex, and was surprisingly independent of the number of neurons in the population (Fig. 3b). Interestingly, these results are consistent with theoretical predictions and anatomical observations in the rat cerebellum [33] and the fly mushroom body [39].

A particularly important quality of the RP models, which is of key biological relevance, is their accuracy in learning a population codebook from a severely under-sampled training set. This would affect how quickly a neural circuit could learn from examples an accurate representation of its inputs. Fig. 3c shows large differences in the performance of pairwise maximum entropy and random projection models (see also SI Fig. S2d), when the sample size is of only a few hundred samples. Pairwise based models (and even more so triplet-based models etc.) fail for small training sets because estimating pairwise correlations with limited samples is extremely noisy when the input neurons are mostly silent. In contrast, the linear summation in the random functions of the RP models means that they are estimated much more reliably with small number of samples (see SI Fig. S3b).

The RP models we presented thus far were trained using standard numerical algorithms based on incremental updates [31], which are non-biological in terms of the available training data and the computations performed during learning. As we demonstrate below, we can find learning rules for RP models that are simple, biologically plausible, and local. While other biologically inspired learning rules may exist, the one we present here is particularly interesting, since noise in the neural circuit is the key feature of its function. Our local learning rule relies on comparison of the activity induced in the circuit by its input 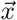 with that induced by a noisy version of the input. This ‘echo’ pattern, 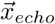, would result from weak and independent noise that may affect each of the input neurons {*x_i_*}, such that 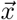 and 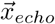 would differ by 1-2 bits on average (see SI for details). Both 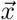 and 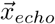 are each propagated by the circuit’s feed-forward connectivity and may result in different activation of the intermediate neurons. If an intermediate neuron is activated only in response to the input but not by the noisy echo, its synapse to the output neuron is strengthened (Fig. 4a); when the converse is true, the synapse is weakened. The updates are scaled by the ratio of the output neuron’s membrane potential *y* in response to the input and its noisy echo. This is concisely summarized in a single learning rule for each of the synapses connecting to the output neuron:

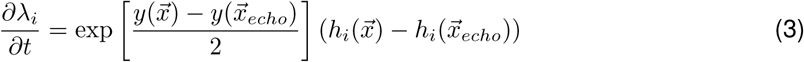

and so the change in synaptic weights depends only on the pre- and post-synaptic activity generated by the most recent input and its echo. This implies that the neural circuit responds with the surprise of its input while simultaneously updating its internal model to account for this input, which also means it can naturally adapt to changing statistics of the input distribution.

**Figure 4:**
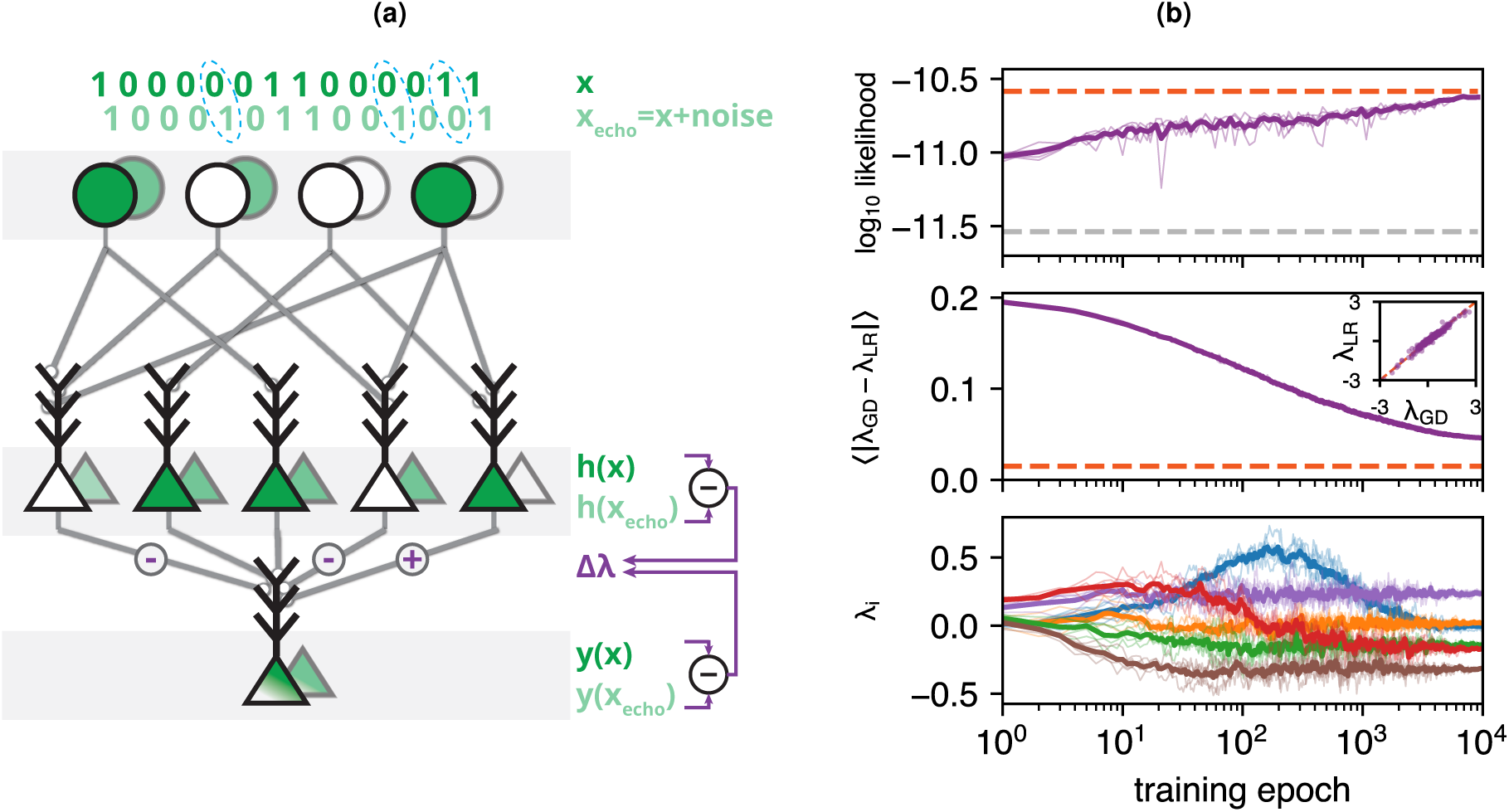
A local biologically plausible learning rule for the RP models based on neural noise gives highly accurate models. **(a)** A learning rule that trains a circuit to respond with the surprise of its input activity patterns by comparing the response to the input pattern (foreground) with the response to a weakly noisy echo of the input pattern (background). Each synaptic weight is modified according to the differences in activity in the pre-synaptic neuron, scaled by the relative membrane potentials of the output neuron. **(b)** RP model trained with the learning rule (LR) and standard gradient descent (GD) on population activity patterns of 100 neurons, by repeatedly presenting epochs of the same 100,000 activity patterns. Bold curves denote average over 5 realizations of learning, each plotted in lighter color. **Top**: Mean log likelihood of test data under the model (dashed orange: model trained with standard GD, dashed grey: independent model). **Middle**: Mean difference in synaptic weights between the learning rule and standard GD (dashed orange: average difference across multiple realizations of standard GD). Inset shows final values of the individual synaptic weights when learned with the learning rule vs. standard GD. **Bottom**: Example of six individual synaptic weights as they are modified across training epochs.

This learning rule induces an average weight change that implements a stochastic gradient descent version of the Minimum Probability Flow [40] algorithm for learning probability distributions (see SI for details and derivation). In this implementation, the neural noise crucially allows the neural circuit to compare the surprise of observed activity patterns with that of unobserved ones, where the goal is to decrease the former and increase the latter. Unlike the traditional role of noise in computational learning theory for avoiding local minima [41, 42] or finding robust perturbation-based solutions [10], here it is the central component that actively drives learning.

Neural circuits trained using the learning rule (Eq. 3) reached a performance close to that of identical circuits (i.e. the same random projections) trained with the non-biological standard gradient descent approach (Fig. 4b top), with closely matching synaptic weights (Fig. 4b middle). These models also accurately captured high-order correlations (SI Fig. S5a) and the distribution of population synchrony (SI Fig. S5b). When trained with severely undersampled data, the performance of RP models trained with the learning rule was comparable to that of the standard pairwise model (SI Fig. S5c).

The RP model can be further improved both in terms of its performance and biological realism by training it using Eq. 3 while periodically discarding projections with a low value of |*λ_i_*| and replacing them with new projections that were selected either randomly (SI Alg. 1) or in such a way that maximizes their predictive contribution (SI Alg. 2). In the equivalent neural circuit, this corresponds to pruning weak synapses to the output neuron (as reported by [43]) and creating new connections to previously-unused parts of the circuit. We found that this simple pruning and replacement of synapses resulted in more compact models, where the performance increases primarily when the model has few projections (Fig. 5a). The pruning, in effect, adapts the random projections to the statistics of the input by retaining those which are more informative in predicting the surprise. Though each intermediate neuron still computes a random function, the set of functions observed after training are no longer drawn from the initial distribution but are biased towards the informative features. As a result, the intermediate units which are retained have lower firing rates and are more decorrelated from each other (Fig. 5b). Thus, when neural circuits learn to compute the surprise of their inputs, pruning weak synapses would result in a more efficient, sparse, and decorrelated activity as a ‘side effect’.

**Figure 5:**
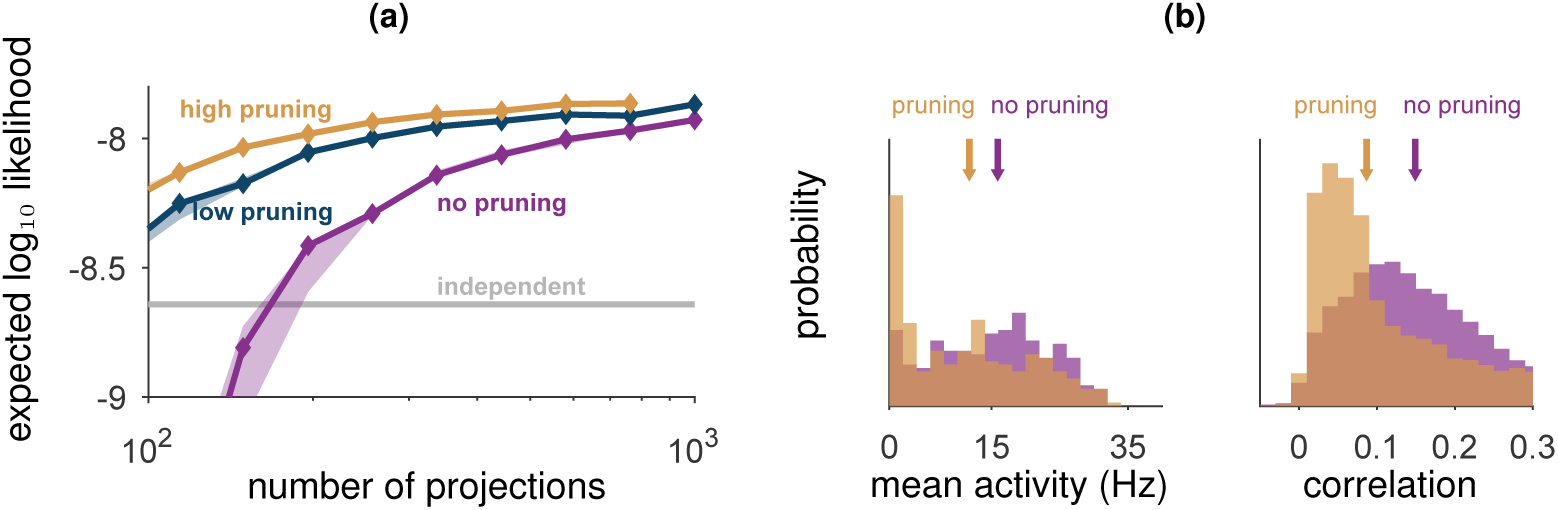
Improved RP models using synaptic pruning and replacement. **(a)** Expected log likelihood of RP models trained with the local learning rule on population activity patterns of 70 neurons from the monkey visual cortex, while periodically pruning weak synapses and replacing them with new randomly chosen projections. Curves denote the performance of models trained with different total average number of replacements per synapse (low: 2, high: 8). **(b)** Average firing rates (left) and Pearson correlations (right) of intermediate units *h_i_* in models trained with the learning rule with (orange) or without (purple) pruning and replacement; arrows denote median values.

## Discussion

The RP models suggest a simple, scalable, efficient and biologically plausible unsupervised building block for neural computation, where a key goal of neural circuits is to generalize from past inputs to estimate the surprise of new inputs. We further presented an autonomous learning mechanism that allows randomly connected feed-forward circuits of spiking neurons to use structure in their inputs to estimate the surprise. These neural circuits can be interpreted as implementing probabilistic models of their inputs that are superior to state-of-the-art probabilistic models of neural codes, while providing greater flexibility and simple scaling to large populations. Our biologically plausible learning rule reweights the connections to an output neuron to maximize the predictive contributions of intermediate neurons, each serving as a random feature detector of the input activity. Relying on noise as a key component, it is a completely local process that operates continuously throughout the circuit’s normal function, and corresponds to a stochastic gradient descent implementation of a known machine learning algorithm. Neural circuits trained this way exhibit various properties similar to those observed in the nervous system: they perform best when sparsely connected and show sparse and decorrelated activity as a side effect of pruning weak synapses.

The estimation of surprise that underlies the RP model also suggests an alternative interpretation to common observations of neural function: feature selectivity of cells would correspond to responding strongly to a stimulus that is surprising based on the background stimulus statistics, and neural adaptation would signify a change in surprise based on the recently observed stimuli. While we focused here on shallow and randomly connected circuits, the local scope of learning in these models also implies they would work in other neural architectures, including deeper networks with multiple layers or networks lacking a traditional layered structure. In particular, this would be compatible with networks where the intermediate connectivity is adjusted by a separate process such as backpropagation in deep neural networks. Importantly, relying on the existing random connectivity as random feature detectors simplifies and accelerates the learning process, and the emerging representations are efficient and sparse [38, 16, 25] without explicitly building this into the model.

The RP model also naturally integrates into Bayesian theories of neural computation: because learning involves only modifying the direct connections to an output neuron, multiple output neurons that receive inputs from the same intermediate layer can each learn a separate model over the stimuli. This could be accomplished if each readout neuron would modify its synapses based on some ‘teaching signal’ only when particular input patterns or conditions occur, thus giving a probabilistic model for new inputs, conditioned on the particular subset of training ones. Thus, comparing the outputs of the readout neurons would give, for example, a Bayes-optimal classifier at the cost of a single extra neuron per input category. More broadly, the RP model would work in concert with general mechanisms of reinforcement learning whose role would be to adapt the firing threshold for the output neuron and provide the gating signal that switches learning on and off according to the external stimuli.

## Online methods

### Experimental Data

We tested our models on extra-cellular recordings from neural populations of the prefrontal and early visual cortices of macaque monkeys. All experimental procedures conformed to the National Institutes of Health Guide for the Care and Use of Laboratory Animals and were approved by the New York University Animal Welfare Committee. For recordings from the visual cortex, we implanted 96-channel microelectrodes arrays (Utah arrays, Blackrock Microsystems, Salt Lake City) on the border of the primary and secondary visual cortices (V1 and V2) of macaque monkeys (macaca nemestrina) such that the electrodes were distributed across the two areas. Recording locations were chosen to yield overlapping receptive fields (RF) with eccentricities around 5° or less. During the experiment, monkeys were anesthetized with Sufentanil Citrate (4-6 µg/kg/hr) and paralyzed with Vecuronium Bromide (Norcuron; 0.1 mg/kg/hr), while drifting sinusoidal gratings were presented monocularly on a CRT monitor [44, 45]. Recordings from the prefrontal cortex were obtained by implantation of 96-channel Utah arrays in the prearcuate gyrus (area 8Ar) of macaque monkeys (macaca mulatta). During the experiments, monkeys performed a direction discrimination task with random dots [46, 47]. Neural spike waveforms were saved online (sampling rate, 30 kHz) and sorted offline (Plexon Inc., Dallas). Throughout the paper we use the term “units” to refer to both well-isolated single neurons and multiunits. Models were fitted in each case to the population activity during all trials, regardless of their difficulty level (for the prefrontal recordings), and over all stimulus induced activity, regardless of the gratings direction or size (in the V1 and V2 data).

### Data Preprocessing

Neural activity patterns were discretized using 20 ms bins. Models were trained on randomly selected subsets of the recorded data, or training set (the number of samples described in each case in the text), and the remaining data was used to evaluate the model performance (held-out test set).

### Construction of random projections

The coefficients *a_i,j_* in the random projections 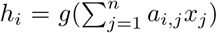 underlying the RP models were randomly set, using a two-step process. First we used a predetermined sparseness value to decide the average number of nonzero values (*indegree*) for each projection, picked them randomly and independently with probability 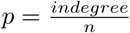(where n is the total number of neuron in the input layer), and set the remaining coefficients to zero. The values of the nonzero elements were then drawn from a Gaussian distribution *a_i,j_* ~ *N*(1, 1). The models were not sensitive to different variants of the selection process of *a_i,j_* (see SI Fig. S3a).

In the results shown in the main text we used *indegree* values in the range of 4-7 (see Fig. 3b for the effect of different indegree values on the model performance) and set *g* to be a threshold function (see SI Fig. S3b for other choices of random functions).

Though the threshold *θ_i_* of each individual projection neuron can be tuned separately, in the results shown in the main text we used a fixed threshold value of 0.1. *indegree* for models trained on the prefrontal cortex and 0.05. *indegree* for models trained on the visual cortex. The models were not sensitive to changes in these values.

### Training probabilistic models with standard gradient descent

We trained the probabilistic models by seeking the parameters *λ_i_* that would minimize the Kullback-Leibler divergence between the model 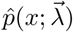 and the empirical distribution *p_emp_(x)*, which is equivalent to maximizing the log likelihood of

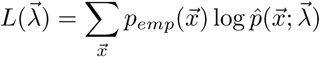

which is a concave function whose gradient is given by

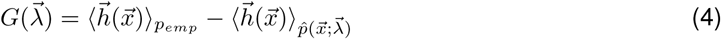

We found the values *λ_i_* that maximize the likelihood by iteratively applying the gradient (Equ. 4) with Nesterov’s accelerated gradient descent algorithm [48]. We computed the empirical expectation in Equ. 4 (left-hand term) by summing over the training data, and the expectation over the parameters 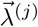 by summing over synthetic data generated from 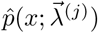 using Metropolis-Hasting sampling.

For each of the empirical marginals 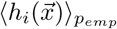we estimated the distribution of marginal values given the empirical one via the Clopper-Pearson method, and assigned it a confidence interval of one standard deviation of this distribution. We set the convergence threshold of the numerical solver such that each of the marginals in the model distribution falls within its corresponding confidence interval. After learning the parameters of the different models, we normalized them using the Wang-Landau algorithm [49] in order to obtain the likelihood of the test data.

We compared the RP model to the independent model, the pairwise maximum entropy model, and the k-pairwise maximum entropy model: The independent model is the maximum entropy model constrained over the mean activities 〈*x_i_*〉, which treats neurons as independent encoders. The pairwise maximum entropy model [17] is the probability distribution with maximum entropy constrained over:

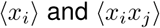

The k-pairwise model [23] uses the same constraints as the pairwise model and adding *n* + 1 syn-chrony constraints:

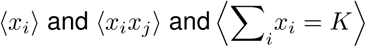

We learned the parameters of the pairwise and k-pairwise models with the same numerical solver used to learn the RP model, and the parameters of independent model by using its closed-form solution. The code used to train the models is publicly available [50] as an open-source MATLAB toolbox: https://orimaoz.github.io/maxent_toolbox/

### MCMC sampling

Synthetic data sampled from the probabilistic models (used in Fig. 2b, S1, S5a and S5b) was generated using Metropolis-Hastings sampling, where the first 10,000 samples were discarded (’burn-in’) and every subsequent 1000th sample was used in order to reduce sample autocorrelations.

### Training RP models with the learning rule

We trained the RP models with the learning rule by iteratively applying the gradient in Eq. 3:

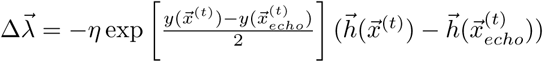

where 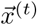 is the joint input to the circuit at time *t*, and 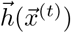 are the concatenated responses of the intermediate neurons *h_1_…h_k_* (see main text). We note that *h* and *y* can be written in vector form using a matrix *A* consisting of the synaptic weights *a_i,j_*:

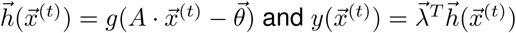

Training was performed over multiple epochs, with the same training data presented on each epoch, and 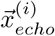 randomly chosen from the training data in each step. The learning rate *η* was set at 0.005 at the first epoch and gradually scaled to 0.00005 in the last epoch, and was normalized by a running average of the gradient norm for numerical stability.

### Training models with synaptic pruning and replacement

To train models with synaptic pruning and replacement, we applied the learning rule with the training data for 10 epochs with decreasing learning rate, and then discarded the 5 projections whose learned values *λ_i_* were closest to zero. We then replaced these discarded projections with new ones either randomly (SI Alg. 1) or in such a way that would maximize the mismatch between the model and the training data (SI Alg. 2). This process was repeated until the desired number of projections were replaced. The performance of these models was not sensitive to different numbers of epochs used or discarded projections.

## Acknowledgements

We thank Udi Karpas, Roy Harpaz, Tal Tamir, Adam Haber, and Amir Bar, for discussions and suggestions, and especially Oren Forkosh and Walter Senn, for invaluable discussions of the learning rule. This work was supported by a European Research Council Grant 311238 (ES), an Israel Science Foundation Grant 1629/12 (ES), research support from Martin Kushner Schnur and Mr. and Mrs. Lawrence Feis (ES), IMH grant R01MH109180 (RS), Simons Collaboration on the Global Brain grant 542997 (RK and ES), Pew Scholarship in Biomedical Sciences (RK), and a CRCNS grant (ES and RK).

## Supporting information

### Self-normalization in RP models

As explained in main text, the membrane potential of the readout neuron 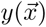 (Eq. 1) reflects the surprise of the input up to an additive factor, which stems from the normalization term of the model (or partition function). In many classification problems, one needs to compare the likelihood values of alternatives, which means only the ratio of probabilities matter and the normalization term cancels out. This would imply that the membrane voltage of the readout neurons would be sufficient. Alternatively, the readout neuron can estimate the normalized value of surprise if we consider an additive term to the membrane voltage that is learned through experience. For example, if the neural code is sparse, the neuron can learn this additive term by taking advantage of the fact that for the all-zero input pattern, 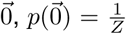 (see [22]) and so the additive factor can be set according to how frequently it receives no spiking input. Yet another alternative would be to consider the spiking of the readout neuron, which would reflect inputs with high surprise, determined by the spiking threshold of the cell. In terms of the spiking of the readout neuron, changing its threshold would be equivalent to an additive term to the membrane potential. As neurons employ homeostatic mechanisms to adjust their activity rates to certain ranges [51], this could be a self-normalizing mechanism for estimating the surprise.

### Other choices of random projection statistics and random function families

RP models were largely unaffected by how the random projection elements were chosen. Fig. S3a shows the cross-validated performance, over experimental data, for models where the synaptic weights *a_i,j_* were drawn from: (1) Normal distribution (*μ* =1, *σ* = 1), (2) Log-normal distribution (*µ* (*μ* =0, *σ* = 1), (3) Uniform distribution in the range [−0.2....0.8] and (4) Binary distribution (*p*(1) = 0.8,p(−1) = 0.2). We found that the different models performed similarly for the data at hand, with the normal and log-normal variants slightly outperforming the others.

We also examined two alternative choices of random function families for a probabilistic model: randomly selected high-order correlations and randomly selected high-order parities. Probabilistic models of randomly selected high-order correlations are maximum entropy distributions constrained over a random selection of high-order correlations, 〈∏*_j,ϵC_i__ x_j_*〉, where *C_1_…C_k_* are randomly chosen groups of neurons in the population. This gives a probabilistic model of the form:

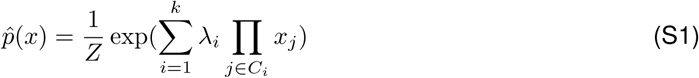

Probabilistic models of randomly selected high-order parities are maximum entropy distributions constrained over the mean parities (XOR) of the activities of randomly selected groups of neurons, 〈 ⊕*_j,ϵC_i__ x_j_*. This gives a probabilistic model of the form:

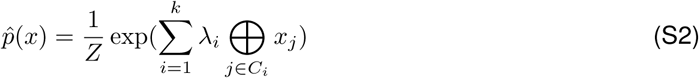

We found that these two models under-performed in comparison to the RP model (Fig. S3b) when trained over our experimental recordings. The particularly poor performance of the correlation-based model can be attributed to the fact that measuring high-order correlations is unreliable when the activity patterns are sparse, as is typically the case in spiking neurons.

### RP models capture high-order interactions

We tested whether RP models can successfully learn data simulated from artificial probability distribution with known high-order interactions by creating a family of parameterized Boltzmann distributions of the form:

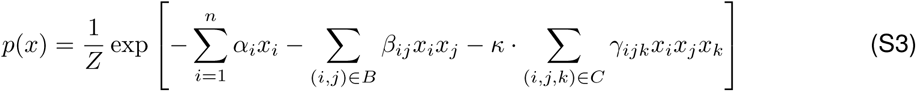

Where *B, C* denote randomly selected pairs and triplets of neurons respectively, and the values*α_i_*, *β_i,j_*, *γ_i,j,k_* were selected randomly from normal distributions. This results in a family of distributions with pairwise interactions when *κ* = 0 and increasingly strong third-order interactions for larger values of *κ*. These distributions were used to generate simulated data, from which RP models were trained. Fig. S4 shows the Jensen-Shannon divergence between the trained models and the original distribution as a function of *I_N_* – *I_2_*, the amount of information in the distribution which cannot be described by pairwise interactions. Maximum entropy models based on pairwise interactions trivially managed to learn when *κ* = 0 but made increasingly large errors as stronger third-order interactions where introduced (larger values of *κ*). RP models were able to capture the high-order interactions more successfully.

RP models trained over population activity patterns of actual neural recordings were able to closely capture high-order correlations in the code. Fig. S1 shows 2nd, 3rd and 4th order correlations in the data and as predicted by RP models trained over a separate training set for neural recordings from the primate visual cortex and PFC.

### Biological interpretation of the ‘echo’ patterns

As explained in the main text, the synaptic learning rule for the RP model (Eq. 3) compares the circuit’s response to the input with its response to a noisy ‘echo’ of the input. Such echo patterns can be generated by biophysical noise either in the neurons or synapses [3]. Updating the synaptic weights to the output neuron would require a backpropagating signal from the cell body [52] and a mechanism that would allow comparing between the synapses’ current and recent activities [53], short-term memory within cells [54, 55], or more complicated local synaptic computations [12]. We note that neural activity during sleep has been characterised with a replay of neural activity statistics alongside highly regular oscillations of neural population activity [56, 57], which could generate replayed inputs with periodically added noisy echos.

### Derivation of the noise-based learning rule from Minimum Probability Flow

Minimum Probability Flow [40] is an algorithm for estimating parameters for probabilistic models by minimizing the KL-divergence between the data and the model after running the dynamics for an infinitesimal time 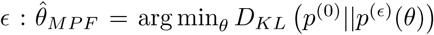. The authors show that this objective function be approximated as minimizing the flow of probability at time *t* = 0, under the dynamics, out of data states *j* ∈ *D* into non-data states *j* ∉ *D* by minimizing the objective:

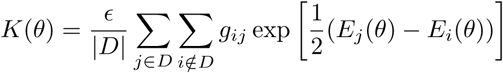

where *g_i,j_* are elements of a binary transition-rate matrix Γ which is allowed to be extremely sparse. This is further extended to the case where Γ is sampled rather than deterministic giving the objective:

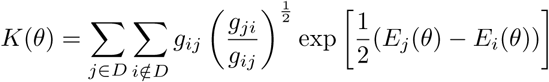

where the inner sum is obtained by averaging over samples from *g_i,j_*. Being a convex function, this term can be minimized by iteratively applying its gradient:

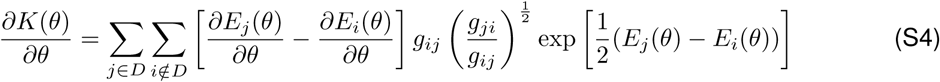

In order to obtain the learning rule we assume that each sample *x* is accompanied by *x_echo_*, a noisy echo of *x* obtained by independently flipping each bit with probability *p* (we typically select *p* such that there is one or two bit flips on average). This is equivalent to setting the following connectivity matrix:

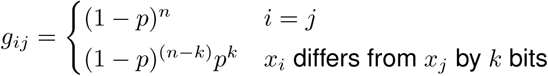

In particular we note that 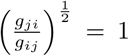 By substituting the model’s energy function *E(x)* = *∑_i_ λ_i_* ⋅ *h_i_* into equation S4 we obtain:

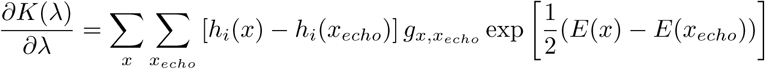

Because the inner sum is obtained by the random generation of *x_echo_* and the outer sum by sampling from the target distribution, a stochastic gradient descent on this gradient would result in the final learning rule:

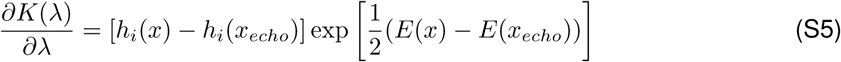

#### Algorithm 1 RP model with pruning and replacement (random selection)

1: Randomly pick a set of *k* projections *h_1_(x)*…*h_k_(x)* for the model

2: Approximately train the RP model on the empirical data *x_emp_*

3: Choose the *q* projections for which |*»_i_*| is smallest and remove them

4: Generate *q* new random projections *g_1_(x)*…*g_q_(x)* and add them to the model

5: repeat steps 2-4 until the required amount of projections has been replaced

#### Algorithm 2 RP model with pruning and replacement (greedy selection)

1: Randomly pick a set of *k* projections *h_1_(x)*…*h_k_(x)* for the model

2: Approximately train the RP model on the empirical data *x_emp_*

3: Choose the *q* projections for which |*»_i_*| is smallest and remove them

4: Generate a set of *r* random projections *g_1_(x)*…*g_q_(x)*

5: Generate a set of samples *x_synth_* ∼ *p*ˆ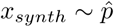

6: Add to the model the *q* projections which maximize: |∑*x_emp_g_j_(x)* – ∑*x_synth_g_j_(x)*|

7: repeat steps 2-6 until the required amount of projections has been replaced

---

**Figure S1:**
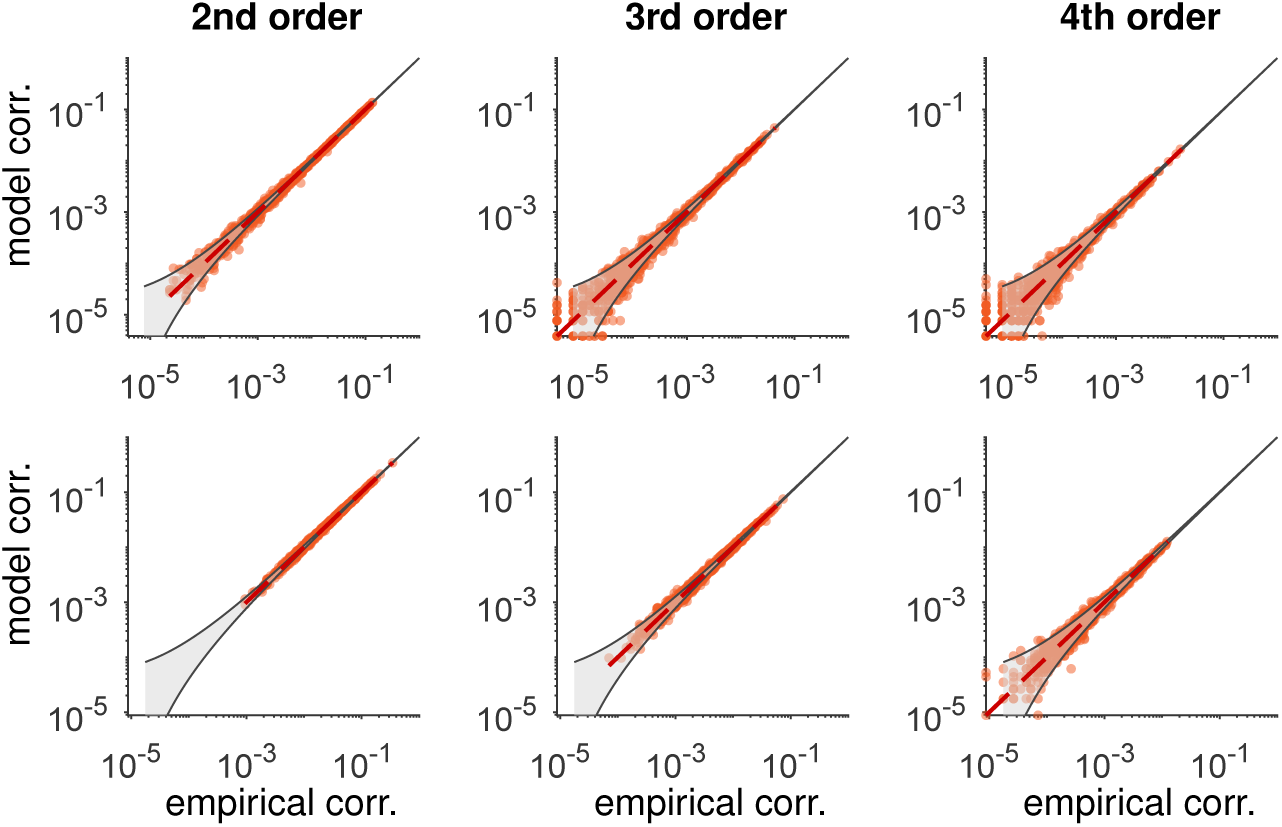
Correlations in test data vs. model prediction for RP models trained over population activity patterns of 178 cells from the visual cortex (top) and 169 cells from prefrontal cortex (bottom).

**Figure S2:**
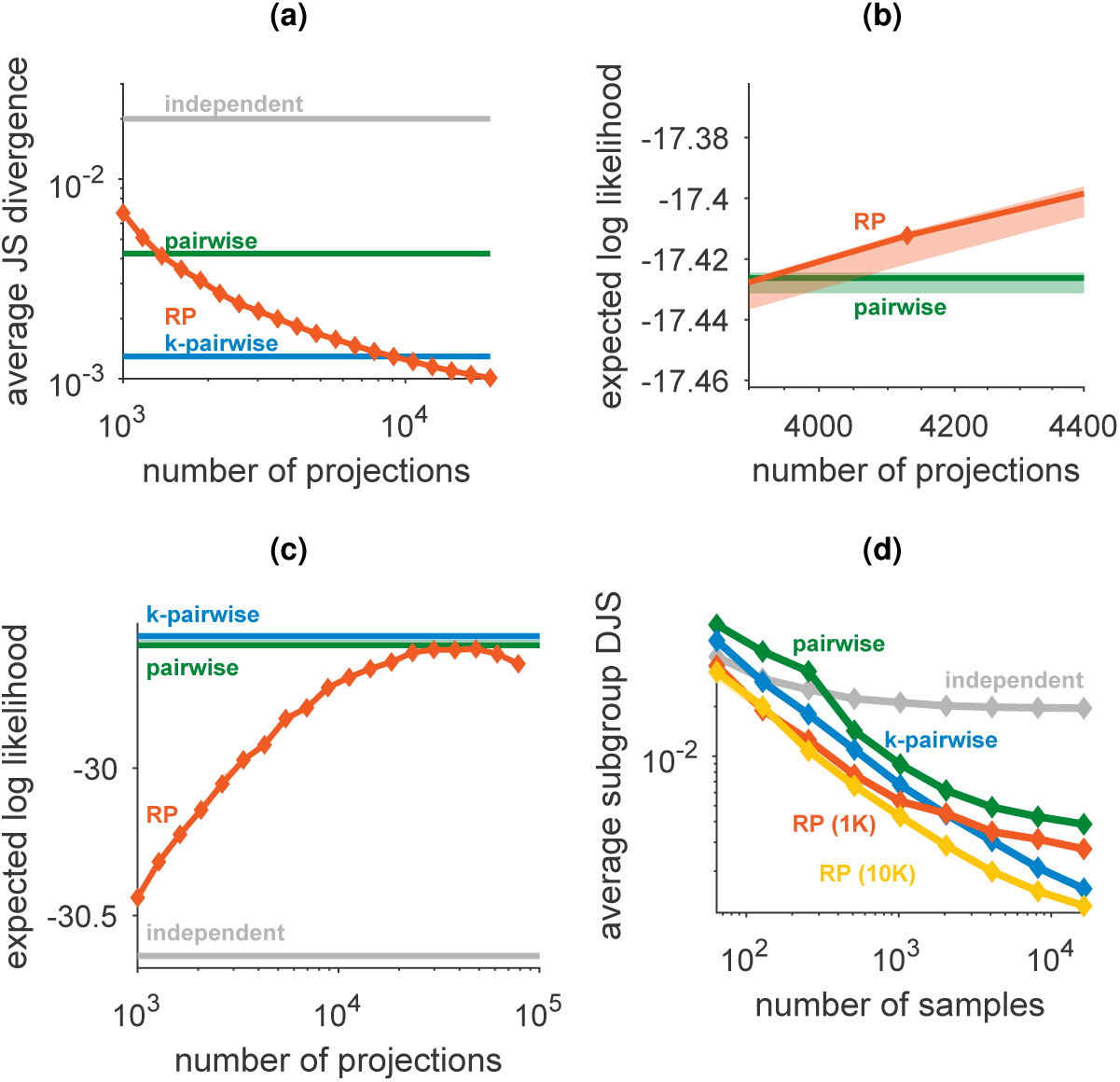
(**a**) Average Jensen-Shannon divergence between model and test data for randomly selected subgroups of 10 neurons. **(b)** Zoomed-in version of figure 3a shows the low variability of RP models for different instantiations of the random projections and random choices of train/test data. **(c)** Expected likelihood of random models trained from population activity patterns from 169 neurons in the prefrontal cortex as a function of the number of projections in the model. The performance of the models, which were trained using 100,000 samples from the population activity, begins to deteriorate as the number of projections approaches this value. Error bars denote standard deviation across random choices of projections and train/test divisions. **(d)** Performance of probabilistic models trained over population activities of 100 neurons from the primate visual cortex, as average Jensen-Shannon divergence between model and test data for randomly selected subgroups of 10 neurons.

**Figure S3:**
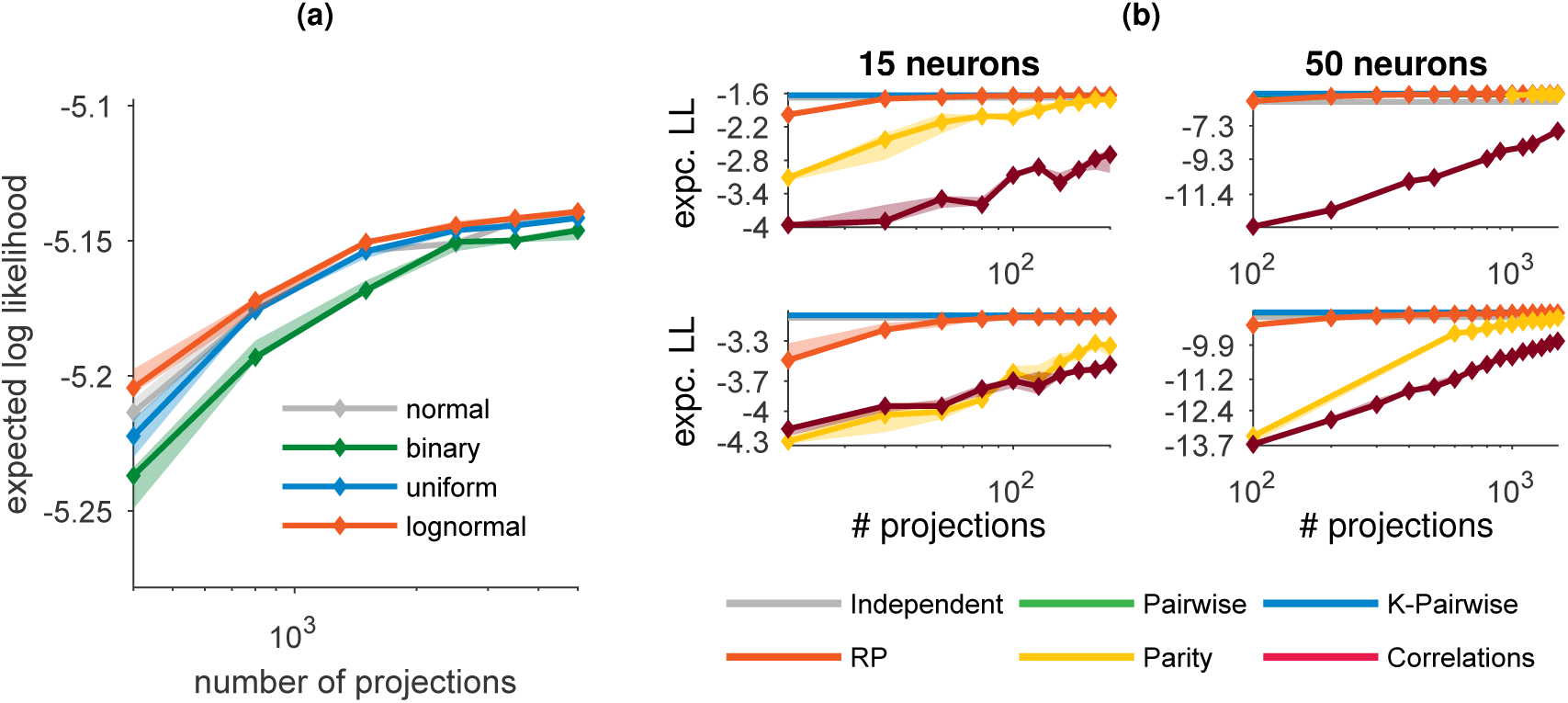
**(a)** RP model performance for different choices of the distributions of the random projection weights. Performance of the RP model compared to models based on high-order correlations and high-order parities of the data, for data from the primate visual cortex (top) or prefrontal cortex (bottom).

**Figure S4:**
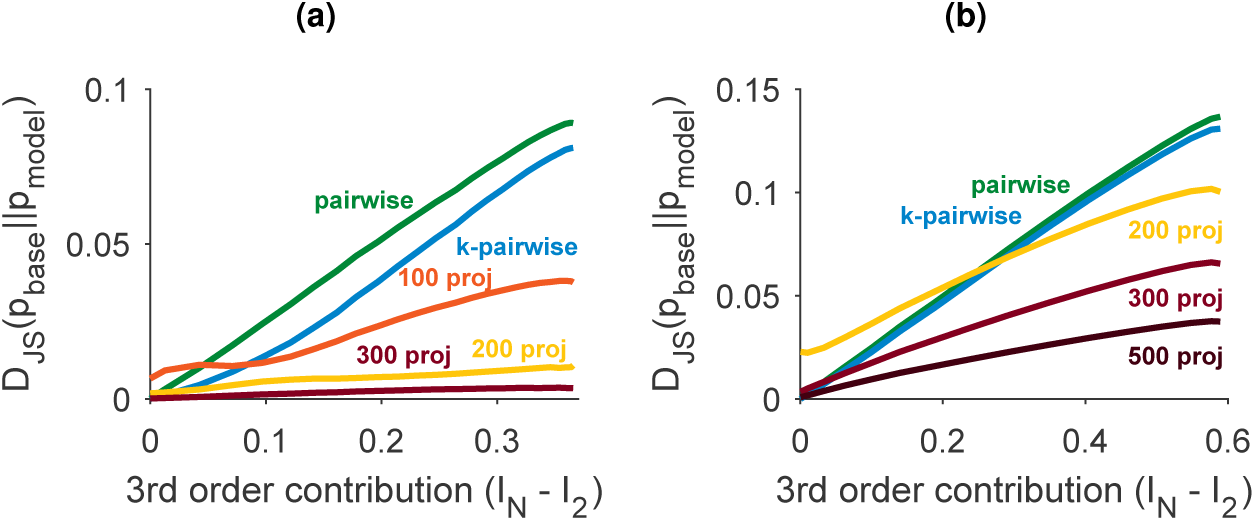
Performance of maximum entropy models as a function of the strength of high-order interactions in small artificial population activity distributions of **(a)** ten and **(b)** fifteen neurons. Pairwise ME models and pairwise with synchrony constraints perfectly captured data generated from pairwise distributions and quickly degraded as stronger third-order interactions were introduced. Random models provided relatively good results even when the third-order interactions were strong.

**Figure S5:**
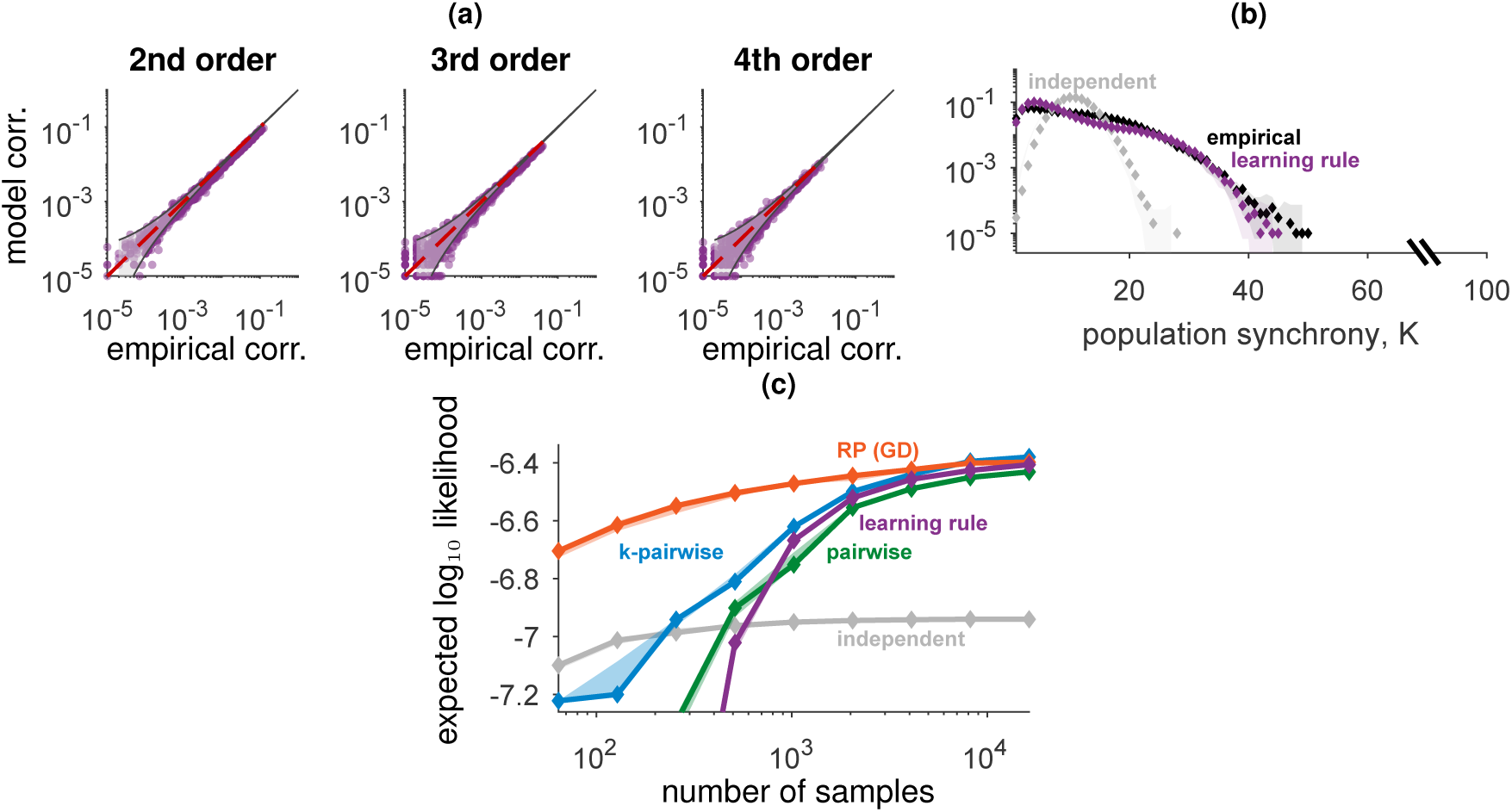
Probabilistic models trained with the biologically plausible learning rule. **(a)** Correlations in test data vs. model prediction for RP models trained with the learning rule over population activities of 100 neurons from the primate visual cortex. **(b)** Population synchrony of RP models trained with the learning rule on groups of 100 neurons, compared to empirical data and reference models. **(c)** Expected likelihood of probabilistic models trained from population activity patterns of 50 neurons from the primate visual cortex as a function of the number of training samples available. In purple: RP model trained with the online learning rule.

